# Preferential activation of the posterior Default-Mode Network with sequentially predictable task switches

**DOI:** 10.1101/2020.07.29.223180

**Authors:** Garazi Araña-Oiarbide, Richard E. Daws, Romy Lorenz, Ines R. Violante, Adam Hampshire

**Affiliations:** Computational, Cognitive and Clinical Neuroscience Laboratory, Department of Medicine, Imperial College London, UK; MRC Cognition and Brain Sciences Unit, University of Cambridge, UK; The Poldrack Lab, Stanford University, US; Department of Neurophysics, Max-Planck Institute for Human Cognitive and Brain Sciences, Germany; School of Psychology, Faculty of Health and Medical Sciences, University of Surrey, UK

**Author notes:** (GAO), (AH).

## Abstract

The default-mode network (DMN) has been primarily associated with internally-directed and self-relevant cognition. This perspective is expanding to recognise its importance in executive behaviours like switching. We investigated the effect different task-switching manipulations have on DMN activation in two studies with novel fMRI paradigms. In the first study, the paradigm manipulated visual discriminability, visuo-perceptual distance and sequential predictability during switching. Increased posterior cingulate/precuneus (PCC/PrCC) activity was evident during switching; critically, this was strongest when the occurrence of the switch was predictable. In the second study, we sought to replicate and further investigate this switch-related effect with a fully factorial design manipulating sequential, spatial and visual-feature predictability. Whole-brain analysis again identified a PCC/PrCC-centred cluster that was more active for sequentially predictable versus unpredictable switches, but not for the other predictability dimensions. We propose PCC/PrCC DMN subregions may play a prominent executive role in mapping the sequential structure of complex tasks.

## Introduction

One of the most consistently replicated findings in the neuroimaging literature is the presence of what is referred to as the default-mode network (DMN), comprising the posterior cingulate cortex /precuneus (PCC/PrCC), ventromedial prefrontal cortex (vmPFC), anteromedial prefrontal cortex (amPFC), inferior parietal lobule (iPL), temporoparietal junction (TPJ), lateral temporal cortex (LTC), temporal pole (TempP), dorsomedial prefrontal cortex (dmPFC), the hippocampal formation (HF) and parahippocampal cortex (PHC) [2,10]. Most commonly, the DMN is evident during resting-state functional magnetic resonance imaging (rsfMRI) as a network comprising brain regions with heightened inter-connectivity [48]. Early evidence from event-related functional MRI (fMRI) studies strongly supported a “task negative” activation profile for the DMN, with the popular interpretation being that this reflected a role in internally generated cognition and disengagement from external demands. More specifically, numerous experiments reported deactivation of the DMN during task relative to rest, and the level of deactivation has been observed to scale with the general cognitive difficulty of the task [26,42,43]. Further evidence has come from studies where higher DMN correlation was observed during rest [22,49]. Furthermore, DMN activity has been reported to increase during tasks that require self-referential processes [2,4,13,57,64].

The last few years have seen a diversification in perspectives on the functional role of the DMN in cognition [11,40]. For example, it has been reported that the DMN reconfigures its internal functional connectivity state when a person is engaged in a task [60–62], and modulates its activity according to experimental demands [37,38,56]. Furthermore, as a task becomes automated through practice (e.g. when applying learned rules), there is a concomitant increase in event-related DMN activity [27, 28, 63]. Collectively, these findings indicate a more active role for the DMN during task performance than was commonly believed, including in learning and executive function.

Most relevantly, a fundamental aspect of human cognitive behaviour is the ability to switch attention between different tasks [18,44]. Switching is considered to be a amongst the most cognitively taxing of tasks, being associated with substantially lengthened behavioural response times (MONSELL). Therefore, it is particualrly notable that activation of some, but not all, sub-regions of the DMN has been reported to increase when switches are performed between perceptually distinct tasks [17,54]. Comparably, heightened DMN activation has been observed during Instruction Based Learning (IBL), when attention is initially oriented towards a new instruction [27] and the rule has to be effortfully mapped into working memory. Notably, in both cases, DMN sub-regions coactivate alongside task positive networks that they characteristically anti-correlate with [11,22,23]. More recent evidence has suggested that the DMN and the multiple demand cortex (MDC), which spans frontal and parietal regions that commonly are activated during challenging cognitive tasks, serve complementary roles in tasks where multi-step decision making takes place [66]. How this fractionation of the DMN and its interaction with task-positive networks relates to the distinct executive sub-processes of switching, remains to be determined.

Here, we examine switching-related brain activity at a fine grain using two novel IBL tasks, which were deployed in separate studies. In the first study (N= 16), we sought to explore the role of the DMN in switching by manipulating the reconfiguration demands along three broad dimensions: (1) the visual-perceptual distance of the switches, (2) the discriminability of the stimuli in the switched to rule set and (3) the predictability of when switches would occur within the sequence of task events. In the second study (N= 16), we examined the relationship between predictability of switching and DMN activity in greater detail. Specifically, the original task was modified to produce a fully factorial design with three predictability dimensions: (1) sequential, when the switch events occur, (2) spatial, where the focus of attention will switch to, and (3) perceptual, what visual dimension the focus of attention will switch to. We tested whether DMN sub-regions were most active when switches were predictable in general for all three dimensions, or whether that relationship was specific to the sequential domain.

## Results

### 0.1 Behavioural results

In brief, the behavioural paradigms consisted of blocks that were designed to manipulate different dimensions of switching demand. Each block started with an instruction slide showing the stimulus-response mapping rules that participants needed to apply, followed by trials where participants needed to match centrally presented probes to the flankers according to the current rule (Figure 1). Trials where a new rule was presented are referred to as Switch trials, and trials where the same rule was applied are referred to as Stay trials (more detailed explanation is provided in the Methods section).

**Figure 1.**
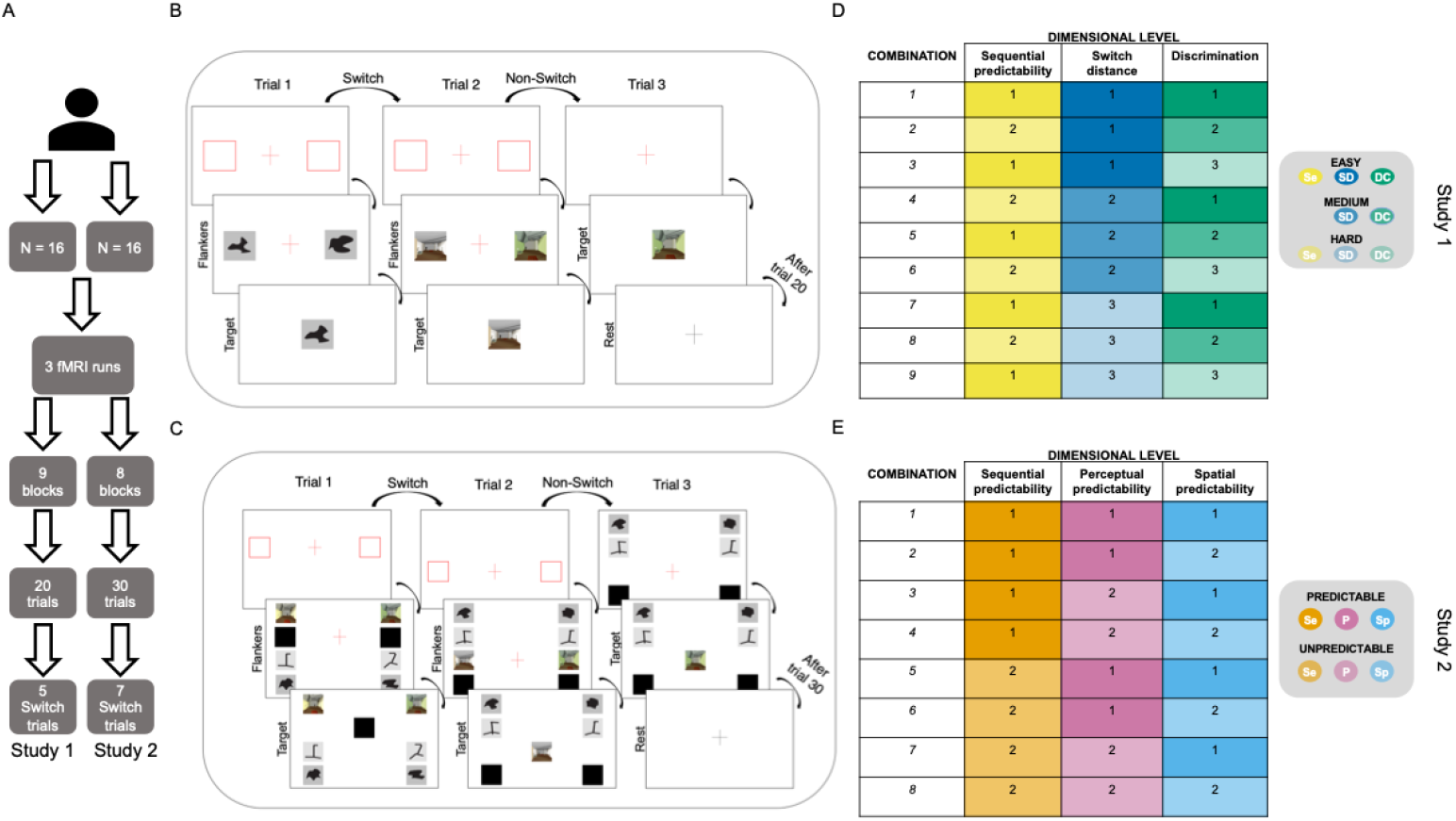
Experimental protocol and cognitive paradigm. **(A)** For each study, 16 participants performed 3 runs in the fMRI scanner. Each run consisted of 9 (Study 1) or 8 (Study 2) blocks of the corresponding task, where each had 20 (Study 1) or 30 (Study 2) trials of which 5 (Study 1) or 7 (Study 2) were switch trials. **(B - C)** On each trial, the participant was first presented with two flankers. They then performed a simple binary-discrimination, where they matched centrally presented probe images to one of the flanker stimuli according to the current rule. **(B)** Study 1. Three aspects of switching were manipulated. (i) The similarity of pre to post switch flanker stimuli (switch distance, SD). (ii) The similarity of the flankers to each other (switch resolution, DC) and (iii) the regularity of when the switch would occur (sequential predictability, Sp). A warning signal (two red rectangles and a red fixation cross) was presented at the start of the task for 500ms, followed by a set of flanker images (4s). After rule presentation, the flankers were replaced by a probe image. The participants had 1500ms to match the probe to the most similar flanker image. **(C)** Study 2. Stimuli predictability was modulated along three dimensions. (i) The frequency of occurrence of switch events (sequential predictability, Se), (ii) the category of the incoming stimuli (perceptual predictability, P) and (iii) the location of the new relevant flanker images (spatial predictability, Sp). In study two, four pairs of flanker images were presented (4s), with the pair that was currently relevant being indicated by the row that the red fixation cross was displayed in. After rule presentation, the relevant flankers were removed and a probe image was displayed where the fixation cross had been. Participants had 1500ms to match the probe the location of the location where the most similar previously flanker had been. In both studies there were two types of trials: Switch trials and non-switch / Stay trials. During switch trials, a new set of flankers was shown to the participants preceded by a warning signal (two red rectangles and a red fixation cross). During non-switch trials a red fixation cross warned the appearance of a new target image (the non-relevant flankers stayed on the screen in Study 2 **(C)**). The face stimuli are replaced here by black squares to avoid potentially identifying information. **(D-E)** The combinations of dimensional levels that were used to define the parameters of each task block in Study 1 **(D)** and Study 2 **(E)**, with the corresponding colour schemes represented to the right. The order of the blocks was randomised for every run and participant.

#### 0.1.1 Study 1

In study 1 we examined the cost of the switch conditions on performance accuracy and reaction time by dividing the trials into Switch and Stay. We computed the accuracy during each of the task blocks, which were designed to vary perceptual distance, discriminability of the probes and sequential predictability of the switches (detailed in Figure 1D). These values were compared using a repeated-measures ANOVA with factors Block and Trial (Switch or Stay). As expected, the mean accuracy level was significantly above chance (p ¡ 0.0001 t-test) (Figure 2A). There was a significant effect of Block [F(8,270) = 2.9.954, p = 0.00349], but not Trial [F(1,270) = 1.117, p = 0.291]. There was no significant interaction effect [F(8,270) = 0.355, p = 0.943]. The significant effect of Block was the result of differences in accuracy between blocks with different levels of target discriminability and visual-perceptual distance of the switches (p ¡ 0.05, paired t-test with Tukey correction). When mean reaction times were analysed using a model of the same design, we observed a main effect of Trial [F(1,270) = 106.906, p ¡ 0.0001], such that Switch trials were significantly slower than Stay trials across all nine blocks (Figure 2B). There was no main effect of Block [F(8,270) = 1.497, p = 0.158] and no significant interaction [F(8,270) = 0.175, p = 0.994]. Therefore, the first paradigm achieved the expected switch cost in reaction times, where switch trials were consistently slower than stay trials, and participants were able to perform the requirements of the task with accuracy that was above chance.

**Figure 2.**
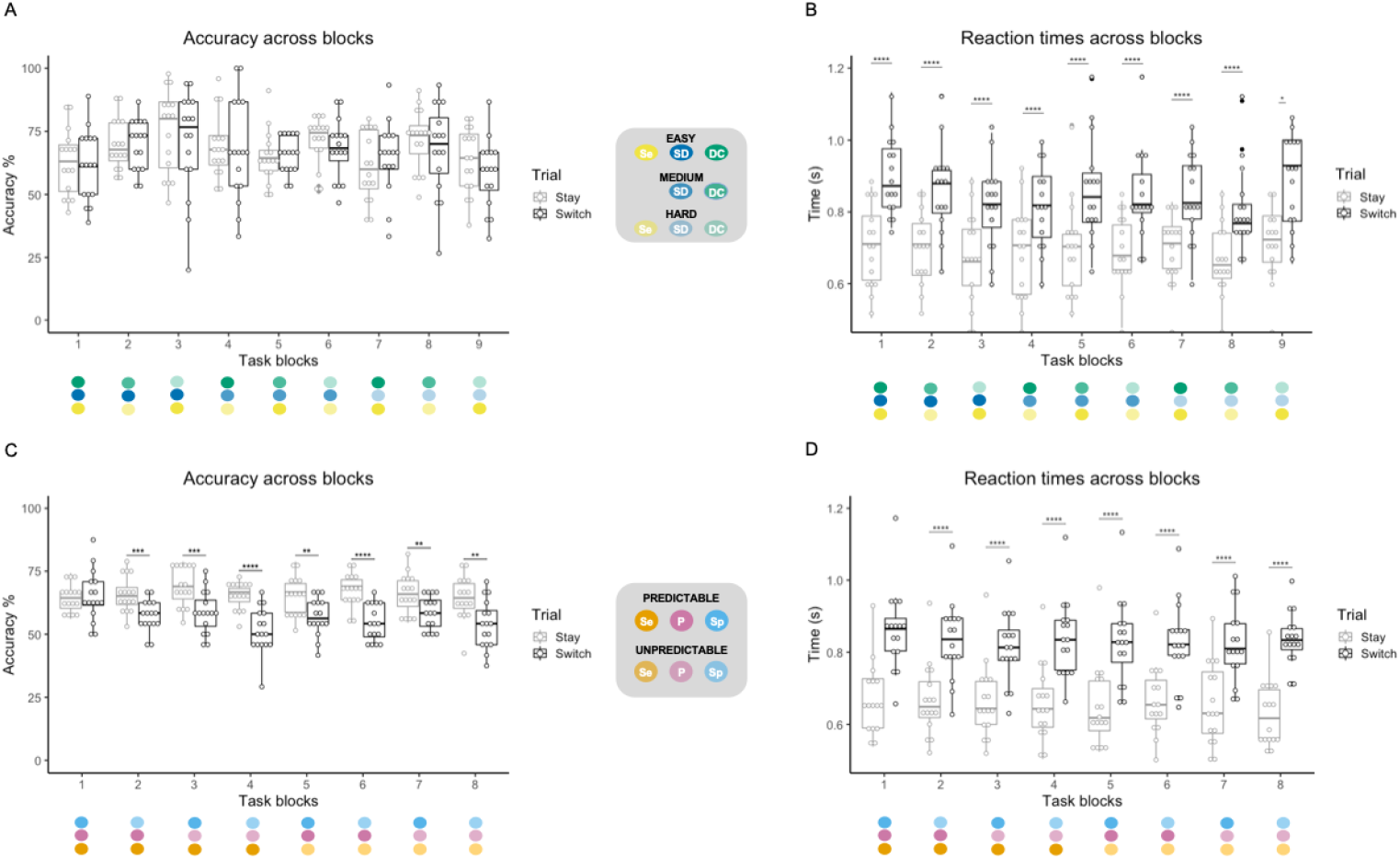
Behavioural results. In Study 1, **(A)** there were no block or trial effects in accuracy (p ≥ 0.05). **(B)** For all nine blocks, switch trials were slower than stay trials (p ≤ 0.0001). In Study 2, **(C)** accuracy was lower for switch trials (p ≤ 0.001), and **(D)** reaction times were slower for switch trials compared with stay trials (p ≤ 0.0001). The coloured spheres in the figure legend refer to the dimensional combinations. Individual participants are indicated by the dots within the plot (n = 16). The top and bottom edges of the boxes represent the 25th and 75th percentiles, with the median being the black line. Extension of the whiskers limits outliers. Significance is indicated by stars (** p < 0.001; *** p < 0.0001), corrected for multiple comparisons with Tukey.

#### 0.1.2 Study 2

The task design in Study 2 was similar to Study 1; however, the stimuli (Figure 10) and the blocks (Figure 1E) were organised in a fully factorial design, capturing all possible combinations of three predictability dimensions. We computed the accuracy during each task block (Figure 1E) for Switch and Stay trials, and compared them using a repeated measures ANOVA with factors for Trial (Switch or Stay) and Predictability (Sequential, Perceptual or Spatial). Accuracy was again significantly above chance (p ¡ 0.0001 t-test) (Figure 2C). There was a significant main effect of Trial [F(1,240) = 83.072, p ¡ 0.001], whereby participants were less accurate in Switch trials relative to Stay trials across all conditions (p ¡ 0.001 t-test with Tukey correction). There was also a significant main effect of Spatial dimension [F(1,240) = 9.685, p = 0.00208], but not the other two dimensions [Sequential F(1,240) = 1.508, p = 0.220; Perceptual F(1,240) = 1.434, p = 0.232]. There were significant interactions between Trial and Spatial, and Trial Type, Sequential and Perceptual factors. Assessment of reaction times across the task blocks for switch and stay trials was performed with a 2×2×2×2 repeated-measures ANOVA for factors capturing Trial (Switch vs Stay) and each of the three predictability dimensions. As intended, there was a significant main effect for the type of trial [F(1,240) = 180.637, p ¡ 0.0001], with the responses for Switch trials being slower than for Stay trials (p ¡ 0.0001) (Figure 2D). There were no significant main effects for the predictability dimensions or the interactions between them. Therefore, the task produced the expected effects on reaction times whilst maintaining a suitable level of accuracy.

### 0.2 Imaging analysis

#### 0.2.1 Posterior DMN areas recruited during switch events

##### 0.2.2 Study 1

Whole-brain voxelwise analysis was conducted to identify clusters of voxels that were active during task blocks relative to rest. 12 clusters were identified in Study 1 for the contrast “Task block > Rest” (Figure 3A left). These were located in the inferior and superior occipital cortices, bilateral frontal pole, superior parietal lobule, pre- and post-central gyri and basal ganglia (peak activation coordinates can be found in Table 1). No DMN regions were observed as active in this contrast.

Conversely, a more widespread activation pattern was observed for task-switching events (“Switch ¿ Rest” contrast, Figure 3A right, Table 2), with a large cluster spanning the occipito-temporal cortex. Notably, the PCC/PrCC region of the DMN was active, but other DMN sub-regions were not. Peak areas outside the DMN included areas of what is often referred to as Multiple Demand Cortex (MDC), comprising the superior and middle frontal gyri and basal ganglia in the left hemisphere, and the right frontal pole, and the supracalcarine cortex and lateral occipital cortex.

**Figure 3.**
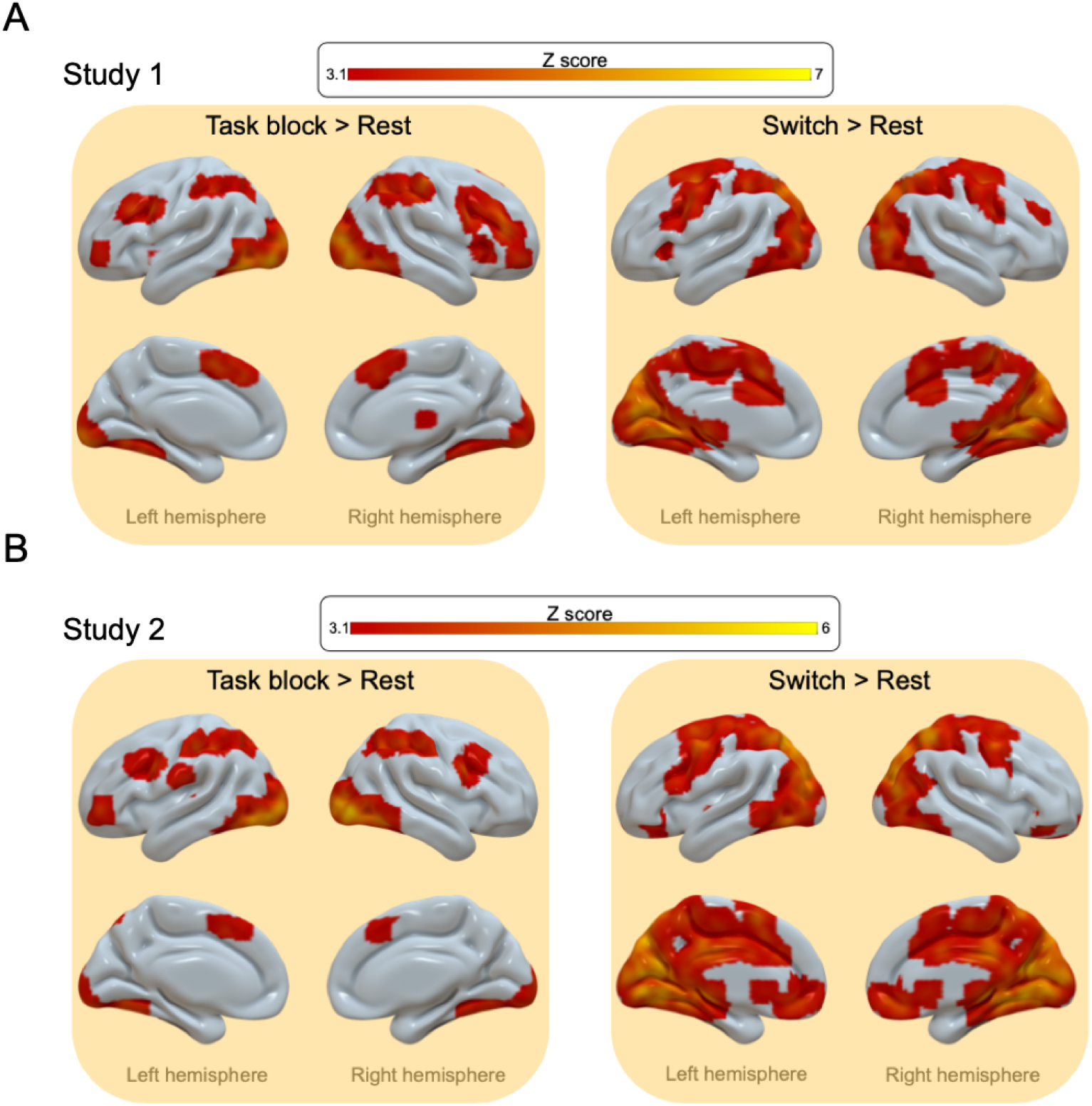
Whole-brain activation during switching task. Activation maps for the contrasts “Task block ¿ Rest” and “Switch > Rest” for Study 1 (**A**) and Study 2 (**B**). Results from the voxelwise analysis were cluster corrected (z > 3.1, p < 0.05) and overlaid in a MNI152 brain template. Peak activation coordinates can be found in Table 1, Table 2, Table 3 and Table 4.

**Table 1.** Results from the cluster-based activation analysis for “Block > Rest contrast” in Study 1. The coordinates indicate the location of the peak activation voxel within a cluster.

| Custer index | Voxels | P | z-max | x | y | z |
| --- | --- | --- | --- | --- | --- | --- |
| 12 | 3942 | 1.75E-27 | 6.2 | 38 | -82 | -2 |
| 11 | 3628 | 5.63E-26 | 6.62 | -20 | -102 | 10 |
| 10 | 1598 | 8.27E-15 | 5.35 | 44 | 36 | 30 |
| 9 | 1387 | 2E-13 | 5.7 | 48 | -28 | 56 |
| 8 | 851 | 1.53E-09 | 4.83 | -44 | -42 | 58 |
| 7 | 600 | 1.79E-07 | 5.28 | -4 | 8 | 52 |
| 6 | 577 | 2.98E-07 | 4.73 | -42 | 8 | 34 |
| 5 | 302 | 0.000164 | 4.22 | -22 | 0 | 14 |
| 4 | 251 | 0.000632 | 5.42 | 46 | 52 | -10 |
| 3 | 227 | 0.00123 | 4.55 | 32 | 26 | 0 |
| 2 | 117 | 0.0376 | 3.91 | -46 | 50 | -6 |
| 1 | 110 | 0.0479 | 4.02 | 16 | -16 | 12 |

**Table 2.** Results from the cluster-based activation analysis for “Switch > Rest contrast” in Study 1. The coordinates indicate the location in MNI space of the peak activation voxel within a cluster.

| Custer index | Voxels | P | z-max | x | y | z |
| --- | --- | --- | --- | --- | --- | --- |
| 4 | 42761 | 0 | 5.81 | 20 | -64 | -2 |
| 3 | 1400 | 4.25E-13 | 4.35 | 28 | 36 | -12 |
| 2 | 157 | 0.0128 | 3.87 | 6 | 18 | -2 |
| 1 | 120 | 4.18E-02 | 4.04 | -30 | 36 | -14 |

##### 0.2.3 Study 2

In Study 2 the contrast of “Task block ¿ Rest” produced similar results. Specifically, we identified 9 clusters (Figure 3B left) within inferior and superior occipital cortices, left-lateralised frontal pole, and pre- and post-central gyri (peak activation coordinates can be found in Table 3). As seen in Study 1, the contrast “Switch ¿ Rest” rendered a broader activation pattern (Figure 3B right, Table 4). The bulk of the activation was within the occipital and temporal regions, with voxels for the DMN bilaterally in the PCC/PrCC. In this occasion other DMN areas also showed activity, specifically the amPFC, PHC and iPL. Outside the DMN, peak activation was located in sub-regions of the MDC, with bilateral clusters around the lateral orbitofrontal cortices (OFC), the lingual gyri and superior lateral occipital cortices. Significant right lateralised activity was also observed within the basal ganglia.

**Table 3.**
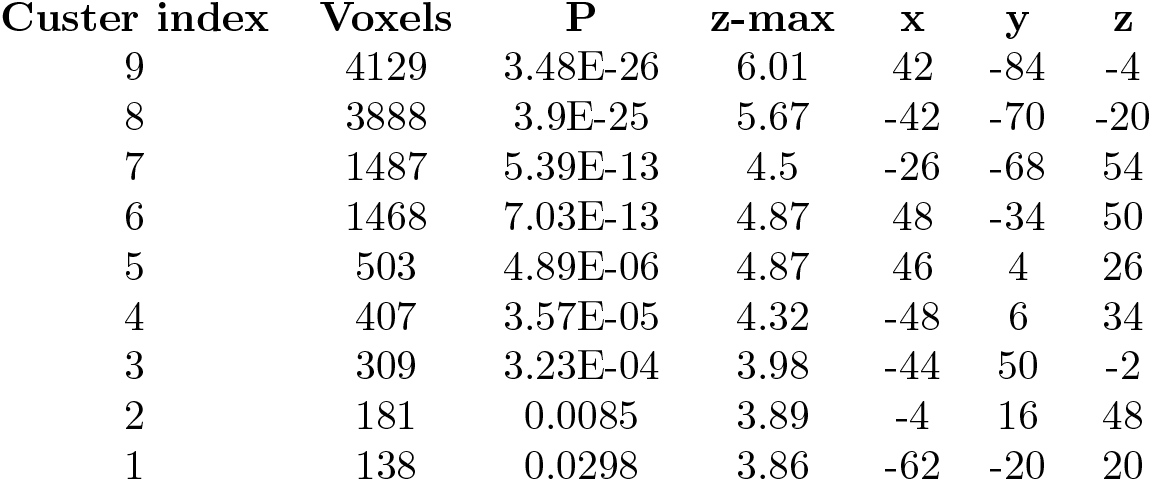
Results from the cluster-based activation analysis for “Block > Rest contrast” in Study 2. The coordinates indicate the location in MNI space of the peak activation voxel within a cluster.

| Custer index | Voxels | P | z-max | x | y | z |
| --- | --- | --- | --- | --- | --- | --- |
| 9 | 4129 | 3.48E-26 | 6.01 | 42 | -84 | -4 |
| 8 | 3888 | 3.9E-25 | 5.67 | -42 | -70 | -20 |
| 7 | 1487 | 5.39E-13 | 4.5 | -26 | -68 | 54 |
| 6 | 1468 | 7.03E-13 | 4.87 | 48 | -34 | 50 |
| 5 | 503 | 4.89E-06 | 4.87 | 46 | 4 | 26 |
| 4 | 407 | 3.57E-05 | 4.32 | -48 | 6 | 34 |
| 3 | 309 | 3.23E-04 | 3.98 | -44 | 50 | -2 |
| 2 | 181 | 0.0085 | 3.89 | -4 | 16 | 48 |
| 1 | 138 | 0.0298 | 3.86 | -62 | -20 | 20 |

**Table 4.**
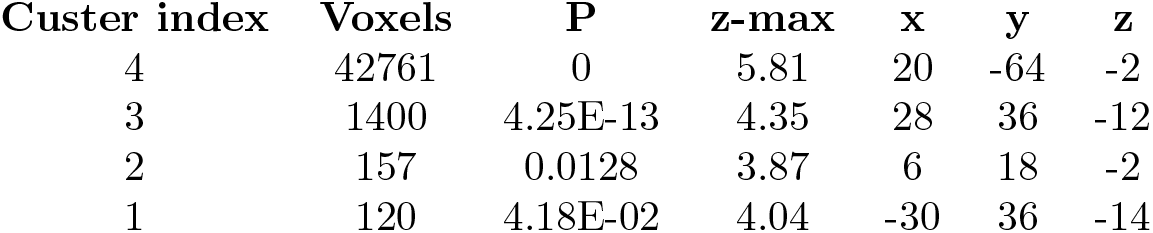
Results from the cluster-based activation analysis for “Switch > Rest contrast” in Study 2. The coordinates indicate the location in MNI space of the peak activation voxel within a cluster.

#### 0.2.4 Preferential activation of the PCC/PrCC to temporally predictable task switches

##### 0.2.5 Study 1

Anatomically distinct regions of interest (ROIs) belonging to the DMN were identified from the “Switch ¿ Rest” activation map. Mean parameter estimates were calculated across all voxels separately for each ROI and each switch condition. These values were examined at the group level using a one-way repeated measures ANOVA for the factor Block. ROI coordinates are in Table 5.

**Table 5.** Regions of interest of Study 1 belonging to the DMN. Coordinates are in MNI space.

| ROI | Hemisphere | x | y | z |
| --- | --- | --- | --- | --- |
| PCC - 1 | L | -12 | -45 | 0.8 |
| PCC - 1 | R | 12 | -45 | 0.8 |
| PCC - 2 | L | -6 | -54 | 8 |
| PCC - 2 | R | 6 | -54 | 8 |
| PrCC - 1 | L | -20 | -60 | 8 |
| PrCC - 1 | R | 20 | -60 | 8 |
| PrCC - 2 | L | -14 | -60 | 8 |
| PrCC - 2 | R | 14 | -60 | 8 |

A statistically robust main effect of Block was evident in the left hemisphere for ROIs in the PCC and PrCC [Figure 4 [PCC F(8,135) = 3.377, p = 0.00146, PrCC F(8,135) = 2.259, p = 0.0269], with no significant effects in the right hemisphere [PCC F(8,135) = 1.844, p = 0.0741, PrCC F(8,135) = 1.315, p = 0.241]. Post hoc tests showed significant block-wise differences between blocks 1 and 5 vs block 6 (p < 0.05, paired t-test corrected with Tukey) and borderline-significant differences between blocks 3 and 6 (p = 0.0643), 4 and 5 (p = 0.0538) and 6 and 7 (p = 0.0626). These blocks were characterised as having different predictability levels (illustrated by the coloured figure legend in Figure 4), where odd blocks had highly predictable switches and even blocks had sequentially unpredictable switches. These results suggest that the activation pattern in both locations was influenced by the level of switch predictability, being higher for more predictable switches.

**Figure 4.**
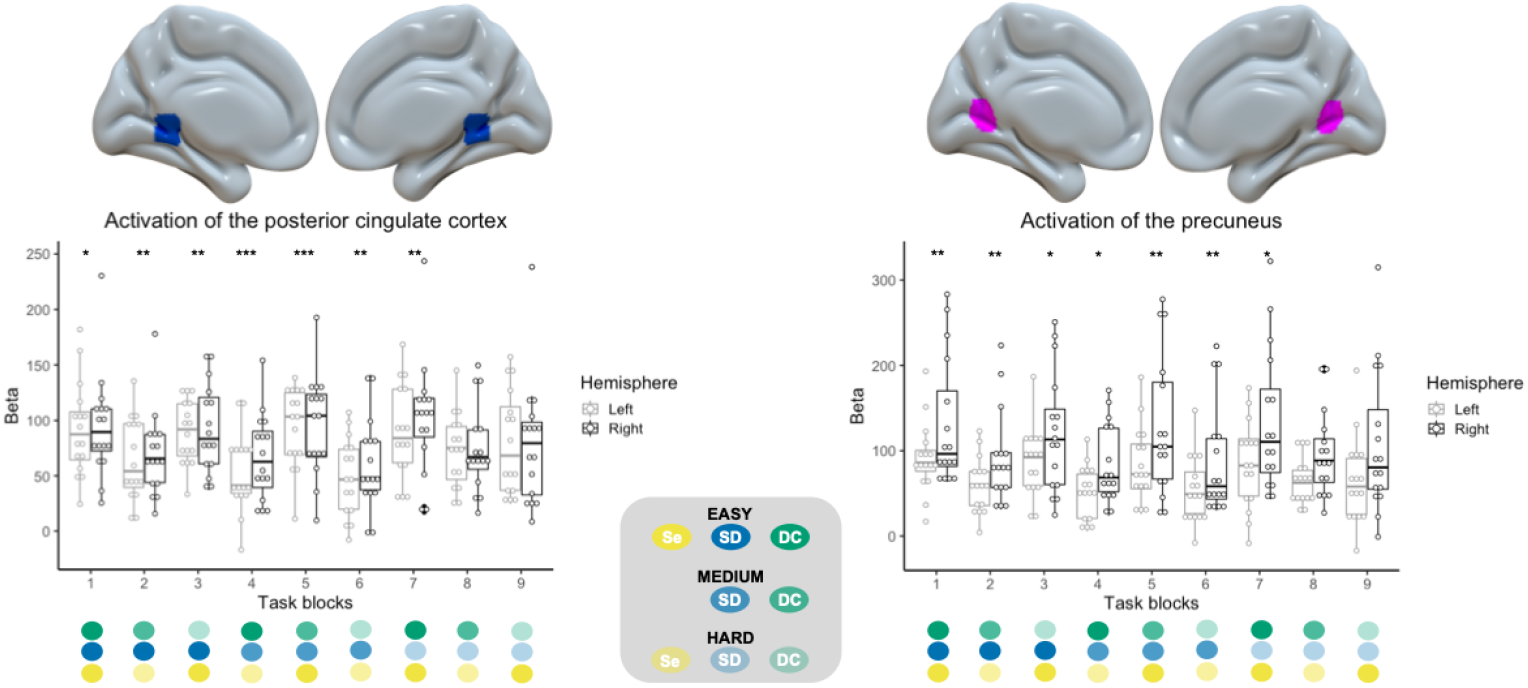
Posterior cingulate/precuneus cortex activation during the switching-task in Study 1. Peak voxel parameter estimates during switch events across the nine task blocks in the posterior cingulate cortex (PCC) and precuneus (PrCC) relative to rest. The coloured spheres in the figure legend refer to the dimensional combinations. Individual participants are indicated by the dots within the plot (n = 16). The top and bottom edges of the boxes represent the 25th and 75th percentiles, with the median being the black line. Extension of the whiskers limits outliers. Significance is indicated by stars (* p < 0.05; ** p < 0.001; *** p < 0.0001), corrected for multiple comparisons with Tukey. The ROIs are overlaid in a MNI152 brain template an and the coordinates can be found in Table 6.

##### 0.2.6 Study 2

In Study 2, we again defined ROIs that belonged to the DMN from the “Switch > Rest” activation map. We extracted mean parameter estimates for each switch condition for each ROI (Table 6 and examined the differences at the group level using a full factorial 2×2×2 repeated measures ANOVA with the 3 predictability dimensions as factors (Sequential, Perceptual and Spatial).

**Table 6.**
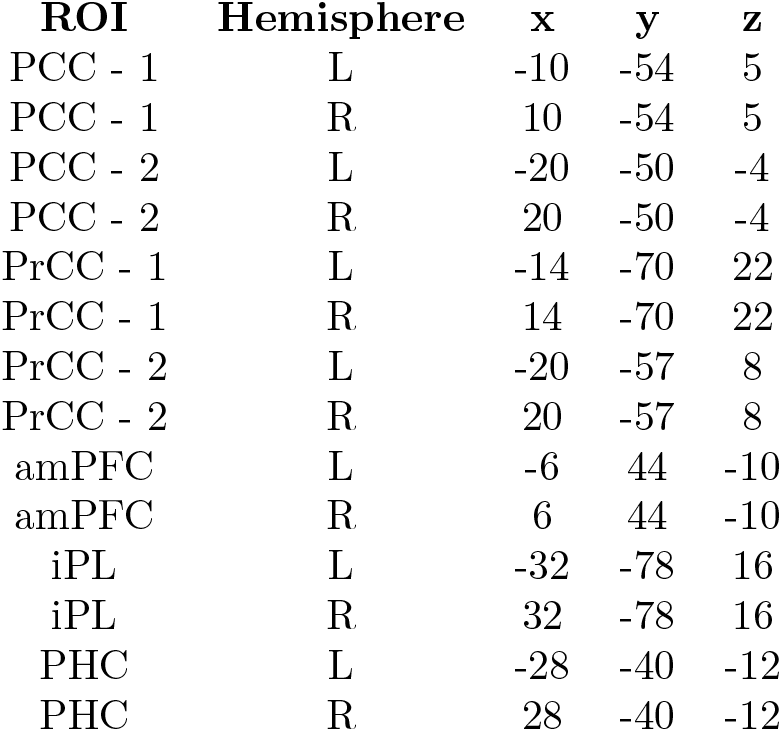
Regions of interest of Study 2 belonging to the DMN. Coordinates are in MNI space.

Confirming the findings from Study 1, there was a significant main effect of Sequential predictability for a bilateral PCC/PrCC set of ROIs [Figure 5, left PCC F(1,120) = 11.282, p = 0.00105, right PCC F(1,120) = 9.246, p = 0.0029; left PrCC F(1,120) = 19.656, p < 0.0001, right PrCC F(1,120) = 10.333, p = 0.00168]. Paired t-tests confirmed higher activation for sequentially predictable switches relative to sequentially unpredictable ones (all p < 0.0001 Tukey corrected). The main effects and interactions for the other predictability dimensions were non significant.

**Figure 5.**
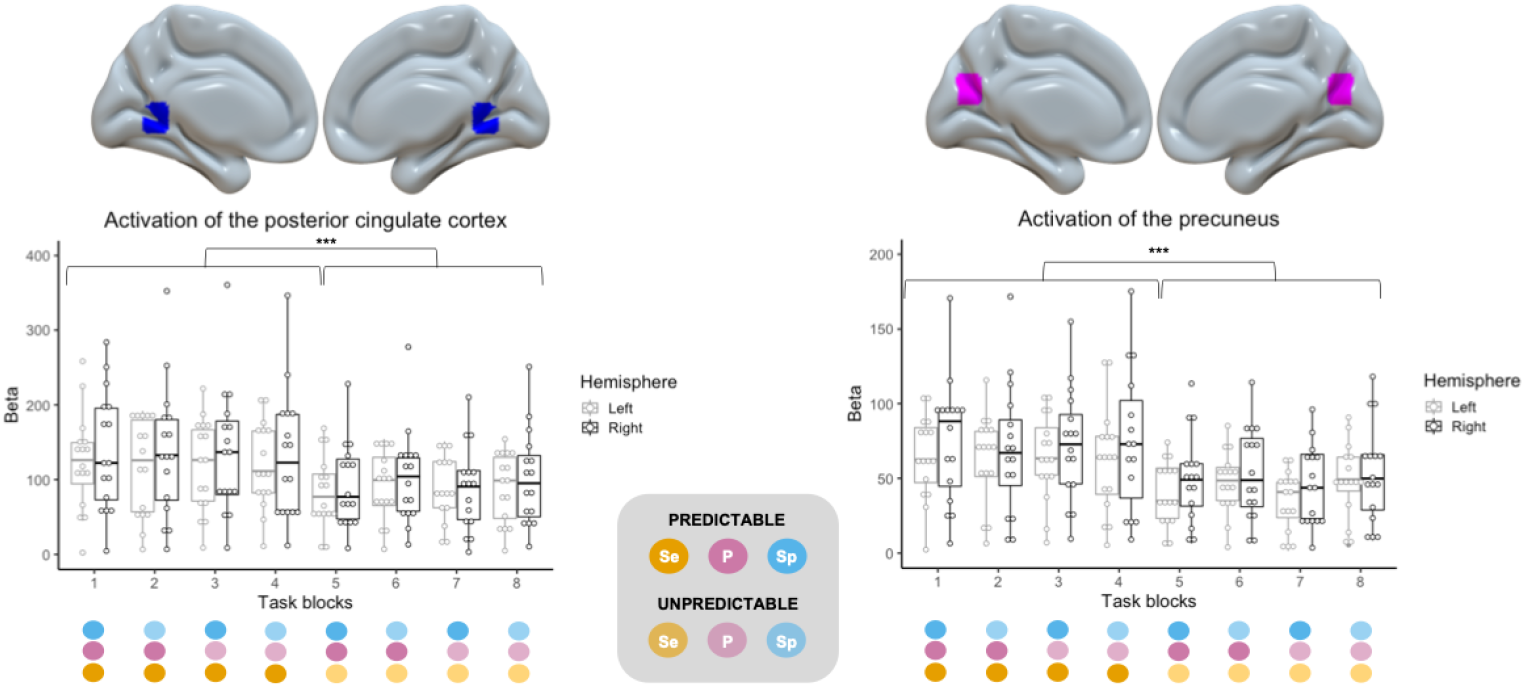
Posterior cingulate/precuneus cortex activation during the switching-task in Study 2. Peak voxel parameter estimates during switch events across the eight task blocks in the posterior cingulate cortex (PCC) and precuneus (PrCC) relative to rest. The coloured spheres in the figure legend refer to the dimensional combinations. Individual participants are indicated by the dots within the plot (n = 16). The top and bottom edges of the boxes represent the 25th and 75th percentiles, with the median being the black line. Extension of the whiskers limits outliers. Significance is indicated by stars (*** p < 0.0001), corrected for multiple comparisons with Tukey. The ROIs are overlaid in a MNI152 brain template and the coordinates can be found in Table 6.

These effects were not changed by placing ROIs in the alternative peak coordinates defined from the “Switch > Rest” contrast in Study 2, with main effects found solely for the Sequential predictability factor [Figure 6, left PCC - 2 F(1,120) = 8.793, p = 0.00365, right PCC - 2 F(1,120) = 21.186, p < 0.0001; left PrCC - 2 F(1,120) = 10.641, p = 0.00144, right PrCC - 2 F(1,120) = 7.571, p = 0.00685]. Post hoc tests showed higher activation related to sequentially predictable switches than unpredictable switches (all p < 0.0001, pair t-test Tukey corrected). Thus, all predictability effects related to the sequential manipulation.

**Figure 6.**
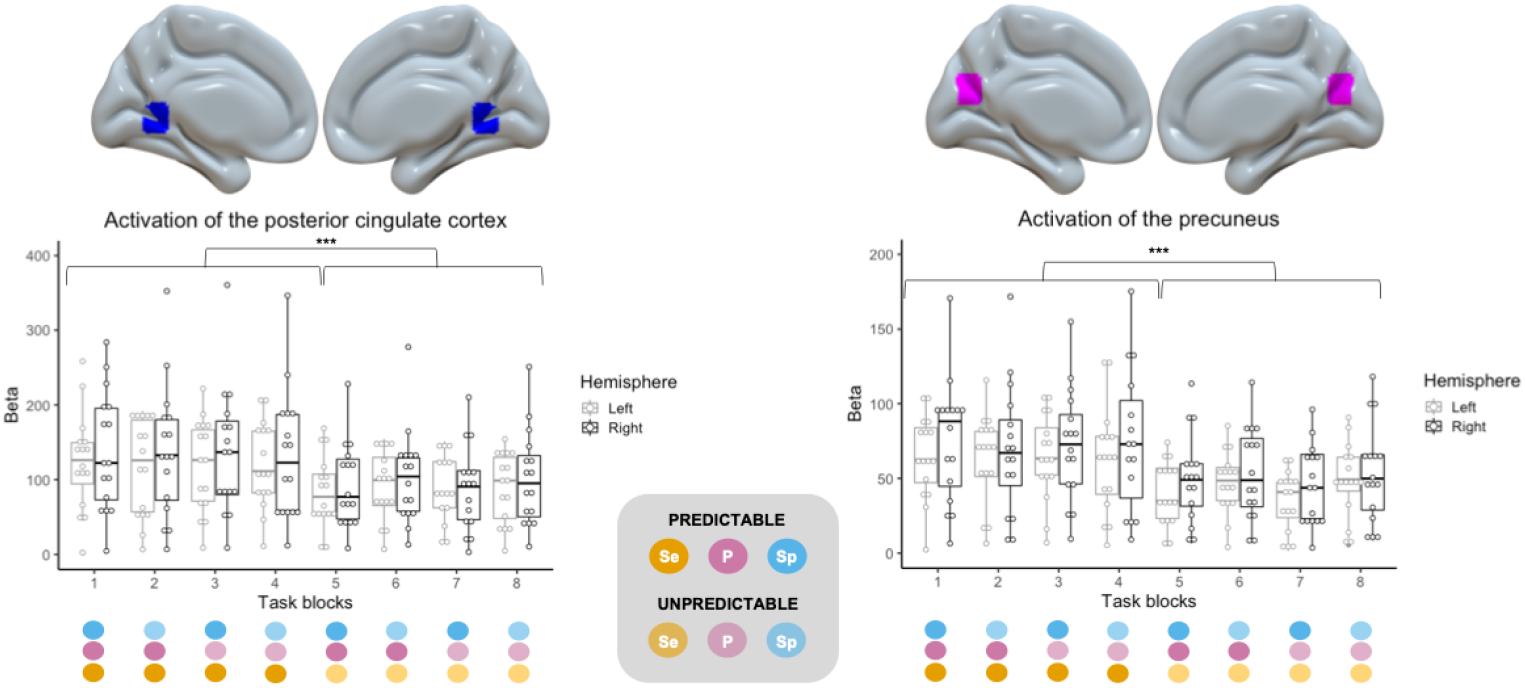
Activation during switch events of a second set of posterior cingulate/precuneus cortex ROIs in Study 2. Activation during switch events of a second set of posterior cingulate/precuneus cortex ROIs in Study 2. Peak voxel parameter estimates during switch events across the eight task blocks in the posterior cingulate cortex (PCC) and precuneus (PrCC) relative to rest. The coloured spheres in the figure legend refer to the dimensional combinations. Individual participants are indicated by the dots within the plot (n = 16). The top and bottom edges of the boxes represent the 25th and 75th percentiles, with the median being the black line. Extension of the whiskers limits outliers. Significance is indicated by stars (*** p < 0.0001), corrected for multiple comparisons with Tukey. The ROIs are overlaid in a MNI152 brain template and the coordinates can be found in Table 6.

To investigate in more detail whether this activation pattern was present in other DMN areas, bilateral ROIs, identified from the “Switch > Rest” contrast of Study 2, in the amPFC, iPL and PHC were examined (Figure 7). No significant main effects were found for the amPFC and the iPL. For the PHC, there was a main effect of Sequential predictability [left hemisphere F(1,120) = 5.418, p = 0.0216; right hemisphere F(1,120) = 4.598, p = 0.034]. As with the PCC/PrCC ROIs, t-tests showed lower activity when switches followed an unpredictable temporal sequence relative to when the sequence was highly predictable (left PCH p = 0.0216, right PCH p = 0.0341 corrected with Tukey). Thus, the manipulation of the sequential structure of the switch events did not only affect the activation of the PCC/PrCC, but also another posterior region of the DMN.

**Figure 7.**
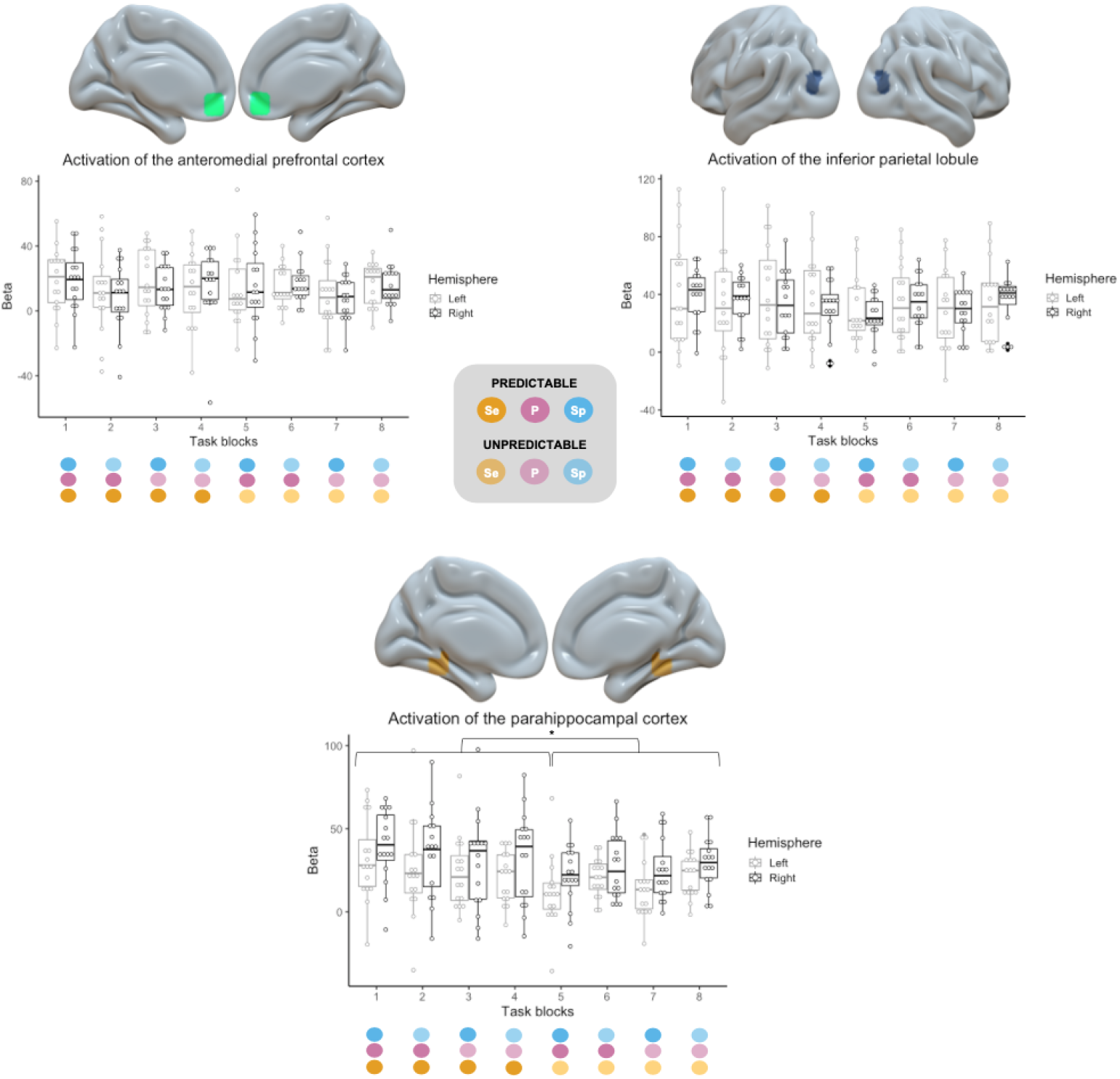
DMN activation during the switching-task in Study 2. Peak voxel parameter estimates during switch events across the eight task blocks in the anteromedial prefrontal cortex (amPFC), inferior parietal lobule (iPL) and parahippocampal cortex (PHC) relative to rest. The coloured spheres in the figure legend refer to the dimensional combinations. Individual participants are indicated by the dots within the plot (n = 16). The top and bottom edges of the boxes represent the 25th and 75th percentiles, with the median being the black line. Extension of the whiskers limits outliers. Significance is indicated by stars (* p < 0.05), corrected for multiple comparisons with Tukey. The ROIs are overlaid in a MNI152 brain template and the coordinates can be found in Table 6.

#### 0.2.7 Sequential predictability, but not perceptual and spatial predictability, modulate posterior DMN acitivity

To support our observations regarding the selective sensitivity of the posterior DMN to sequentially predictable switches we performed an additional analysis in Study 2. A group-level model was constructed to anatomically segregate the effects of each predictability dimension during switch events (whole-brain analysis, cluster activation threshold of z > 3.1 and significance of p < 0.05, Gaussian-Random Field corrected). For the sequentially predictable switches, two significant activation clusters were identified for the contrast “Sequentially predictable > Unpredictable”. The peak activation areas were localised to the right PCC and left PrCC (Figure 8A), in accordance with the results from ROI analysis. In contrast, neither the perceptual predictability nor the spatial predictability (Figure 8B) of the switches showed any significant clusters of brain activation at the corrected threshold.

**Figure 8.**
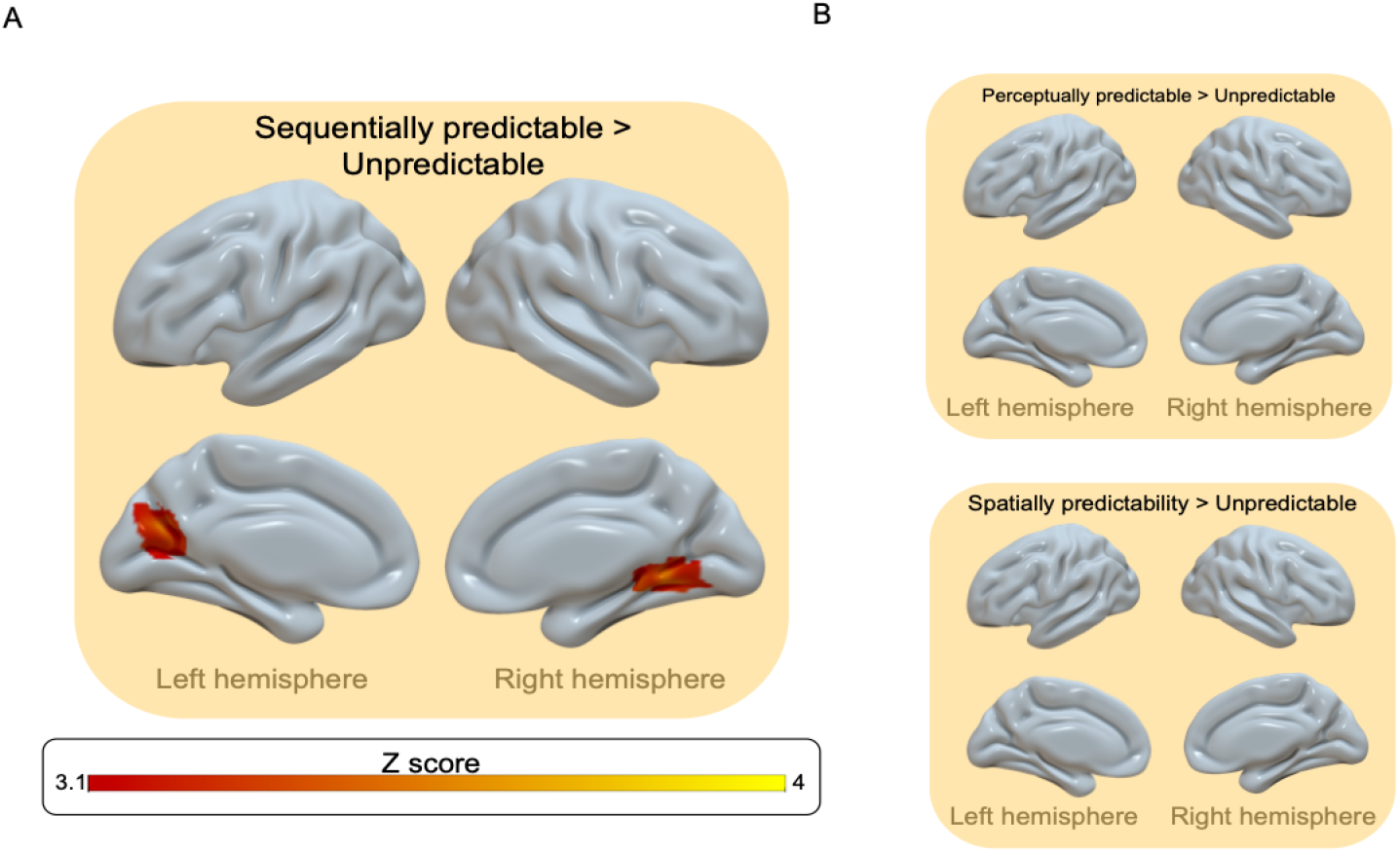
Brain activation during the switching task. Activation maps for the contrasts (A) “Sequentially predictable > Unpredictable”, (B) “Perceptually predictable > Unpredictable” and “Spatially predictable > Unpredictable”. Results from the voxelwise analysis were cluster corrected (z > 3.1, p < 0.05) and overlaid in a MNI152 brain template. Peak activation coordinates can be found in Table 7.

**Table 7.** Results from the cluster-based dimension activation analysis. The coordinates indicate the location in MNI space of the peak activation voxel within a cluster.

| Custer index | Voxels | P | z-max | x | y | z |
| --- | --- | --- | --- | --- | --- | --- |
| 2 | 202 | 0.000194 | 3.97 | 20 | -50 | -4 |
| 1 | 160 | 0.00738 | 3.76 | -14 | -70 | 22 |

To further investigate if there were sub-threshold effects of perceptual and spatial predictability, we performed ROI analysis utilising four active clusters (voxel-activation threshold of z > 3.1) identified in the “Sequentially predictable > Unpredictable” contrast as masks. No significant differences in activation were detected for the contrasts “Perceptually predictable > Unpredictable” and “Spatially predictable > Unpredictable” in any of the four clusters (Figure 9).

**Figure 9.**
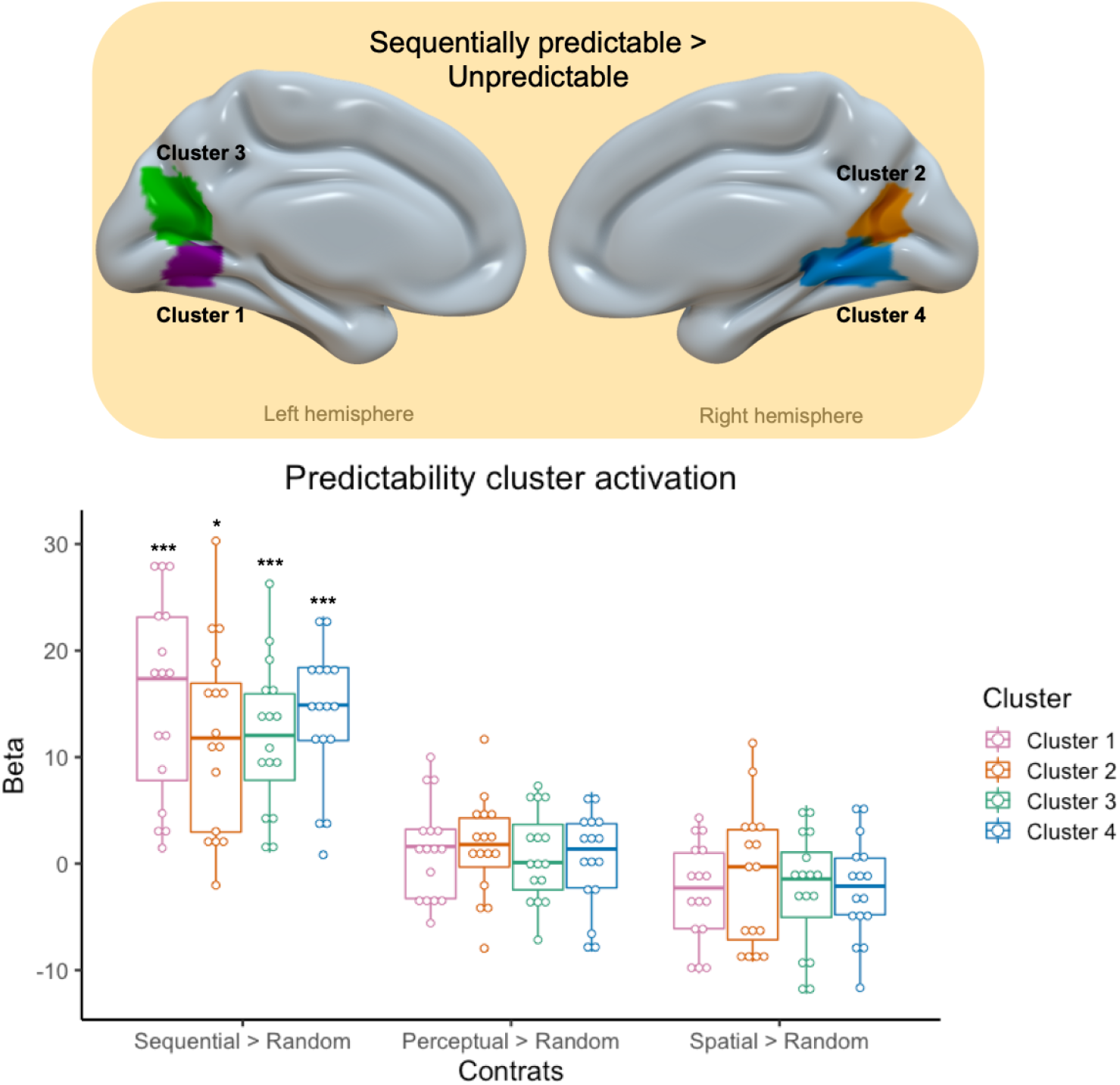
Mean ROI activation across the three predictability dimensions. The group activation map from the contrast “Sequentially predictable > Unpredictable” was used to create brain masks for each of the active ROIs (z > 3.1, >10 voxels). The resulting masks are overlaid in a MNI152 brain template. Peak activation coordinates can be found in Table 8. Mean peak voxel parameter estimates from the four clusters for the contrasts “Sequentially predictable > Unpredictable”, “Perceptually predictable > Unpredictable” and “Spatially predictable > Unpredictable” (cluster corrected at z > 3.1, p < 0.05). Individual participants are indicated by the coloured dots within the plot (n = 16). The top and bottom edges of the boxes represent the 25th and 75th percentiles, with the median being the bold line. Extension of the whiskers limits outliers. Significance after Bonferroni correction is indicated by stars (* p < 0.05; *** p < 0.0001).

**Table 8.**
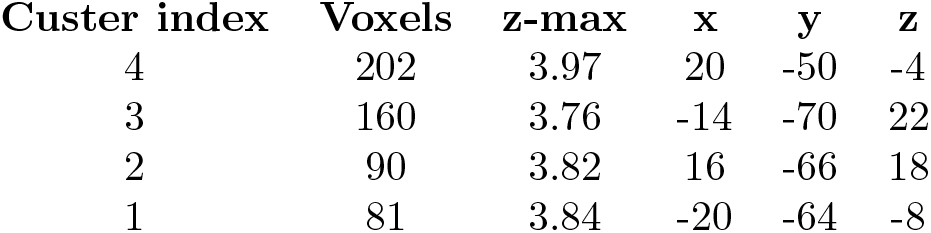
Active clusters from the “Sequentially predictable ¿ Unpredictable” contrast. Clusters were formed with a voxel activation threshold of z > 3.1, and clusters containing > 10 voxels selected. The coordinates indicate the location in MNI space of the peak activation voxel within a cluster.

| Custer index | Voxels | z-max | x | y | z |
| --- | --- | --- | --- | --- | --- |
| 4 | 202 | 3.97 | 20 | -50 | -4 |
| 3 | 160 | 3.76 | -14 | -70 | 22 |
| 2 | 90 | 3.82 | 16 | -66 | 18 |
| 1 | 81 | 3.84 | -20 | -64 | -8 |

## Discussion

We observed a pronounced sensitivity of the posterior DMN to task-switching events, particularly those that follow a sequentially-predictable pattern. This increase in switch-related activation was not present in response to spatially predictable events or those that involved switching to predictable visual dimensions, but it was reproducible across two different tasks. Importantly, this anatomically focused activation was not a characteristic of switching in general, which recruited a much broader distributed set of brain regions.

The preferential activation of a PCC/PrCC cluster with sequentially predictable switches highlights the complexity of cognitive control processes, and has implications for our understanding of the functional role of DMN sub-regions in cognition [11,40]. Imaging studies have reported the involvement of the PCC/PrCC when temporal regularities occur during experimental tasks [1,12] and in instances when actions become repetitive [63]. Our results compliment those and provide an additional facet by showing transient heightened activation was stronger when changes in task rules followed a predictable sequence of events. One possible interpretation is that the PCC/PrCC may code a broader task-representation that does not only include the current set of task rules, but also when this task-set is going to change across time. This would fit with [5]’s account of the proactive brain which states that the brain is able to associate memory representations with the current surrounding and anticipate context-specific aspects of the environment based on predictions that are drawn from these associations.

It could alternatively be argued that the results observed in these two studies were solely caused by the requirement to retrieve information from memory, as reported in previous studies [36,45,51,56]. It is not clear though why predictable switches would involve greater memory retrieval than unpredictable switches. Furthermore, even if this were the case, we would still expect to see significant DMN activation when contrasting the whole task block relative to rest, because both trial types (switch and stay) required the processing of memorised information. The lack of such activation pattern suggests the effect was due to the switch in rules. Although this had a memory component, it was the presence of a task switch that was linked with the signal elevation. Indeed, there is recent evidence implicating the DMN in task-switching [17, 54]. More specifically, those two studies demonstrated a fractionation in DMN when large perceptual switches occurred in the task, with the activation of the PCC/PrCC, PHC, vmPFC and amPFC co-occurring with that sub-regions of the multiple-demand network. Network re-configuration has also been observed in IBL, where presentation of new rules results in global signal elevation across distributed brain regions including the DMN [27, 55], and during a multi-step task with the initiation of a new episode [66]. This may be indicative of an increase in resource-allocation in response to increase cognitive demands.

Regarding the posterior DMN, the relevance of the PCC/PrCC in detecting rule switches accords with results from animal studies of cognitive-set shifting [47]. Single-neuron recordings from these regions in rhesus macaques have shown suppression of activity when they had to switch tasks guided by an explicit signal [31], as well as when the animals were engaged on a task [30]. Conversely, when the animals had to perform a dynamic foraging task without exogenous cues, increases in neuronal firing predicted behavioural shifts [46]. Internally guided and predictable switches may be considered analogous, as both afford the opportunity to internally prepare for the switch occurring at a particular point in time. Thus, the results from guided switch condition could be explained by the lack of requirement to conduct internal evaluation, whereas the formation of an internal model to guide or predict task switches may be evident as increased neuronal firing [47].

Taking a more holistic view, theoretical models of the DMN have argued that this distributed network is composed of functionally dissociable anatomical regions [2,3,19,67]. In concert, they are particularly well positioned to monitor and predict environmental changes. There, the PCC/PrCC has been highlighted as sub-serving a change-detection role in an internally-driven manner [5, 20, 47, 58]. The current empirical results are compatible with that perspective, insofar as the participants monitor, learn and predict the sequential structure of switch events.

Complementing the views discussed so far, episodic memory research has revealed the role of the hippocampus and the entorhinal cortex in temporal sequence learning (for a comprehensive review see [8]), with the entorhinal cortex encoding the temporal structure of an episode [7]. The HF and the PHC are considered part of the DMN, with distinct connectivity profiles at different stages of memory formation [32]. The PHC has been shown to mediate the connectivity between the PCC and the hippocampus [65]. Event boundaries are crucial variables for sequential memory formation and recall [21,25]. The switch trials of our two tasks can be considered event boundaries, in the way they separate the tasks into different sections, and we found some sequential-predictability effects in the activation of the PHC. Future work could expand on this by investigating whether functional connectivity between the hippocampus, the entorhinal cortex, the PHC and the PCC/PrCC is modulated by sequential task-switches.

Additionally, the widespread activation observed in our two studies demonstrates the flexible nature of network membership. We have observed activation in some DMN sub-regions but not others during switching, and this coincides with activation of structures with which the DMN often anti-correlates. Can the label ‘DMN’ be used in these circumstances? It seems more accurate to state that the nodes from which the DMN is comprised can also be members of other functional networks. For the PCC/PrCC this includes a transient conjunction with frontoparietal networks during switching.

If that is the case, then functional networks are highly flexible in their topology and membership, where demands imposed by the task at hand modulate the functional architecture of the brain away from the steady state dynamics of the resting state architecture in order to support particular processes. This accords with the notion of a many-to-many functional mapping between cognitive processes and functional regions of the brain [39, 55].

A caveat when comparing our results to studies of switching is that the two tasks used here did not use classic task-switching designs. Instead, they built on more contemporary instruction-based learning paradigms [14–16, 27, 28, 50]. These differ insofar as the switching of rules is driven by the presentation of an instruction slide with explicit pictorial depiction of the rules, i.e., as opposed to some cue to switch between alternative mappings that are coded within working memory. This type of design is analogous in several key ways. The task program must be updated in response to the instruction cue, and the reduction in RT on trials after new rules were presented relative to when rules remained the same is a characteristic feature of switching. It is notable though that the sequentially predictable and unpredictable switches did not differ with respect to the behavioural switch costs; this is useful because it mitigates the possibility that activation differences between those conditions relate in a trivial way to general difficulty or stimulus processing time. A further possible caveat of our study could be the simple experimental design that makes inferences to real-world situations challenging. However, the simple design allowed disentangling the effects of different task dimensions, and it is a necessary first step to better understand the role of the DMN and its different sub-components in task switching. A straightforward next step is to test whether our results hold in a naturalistic switching task (e.g. movie viewing or virtual reality) where real-world relevance could be more directly inferred. Indeed, the ability to learn and predict the sequential structure of events likely has substantial importance in our complex daily lives. Future work may examine whether disruption of the network dynamics associated with sequential predictability is evident in clinical populations who are prone to executive deficits.

In summary, we have provided evidence for the engagement of the DMN in two task-switching experiments where the implicit sequential structure of switch events was accompanied by the heightened activity of a PCC/PrCC cluster. These results accord with recent accounts of the DMN playing a pivotal role in executive function by utilising internal event representations.

## Materials and Methods

### 0.3 Participants

For the first part of the study we recruited 16 healthy volunteers (12 females, mean age 24.13 ± 4.30 standard deviation, 1 left handed). 9 of those participants also took part in the second study, which had an additional 7 volunteers to bring the total to 16 participants (10 females, mean age 24.47 ± 4.63 standard deviation, 1 left-handed). Participants reported no history of neurological or psychiatric conditions and had normal or corrected vision. The Hammersmith and Queen Charlotte’s Research Ethics committee approved the studies, which conform with the Declaration of Helsinki. All participants gave their written consent after being informed about the nature of the study.

### 0.4 Task switching paradigm

The two switching tasks were programmed in [41] using Psychophysics Toolbox extension [9], based on a previously-reported simple instruction-based learning paradigm [27, 28]. In short, participants had to perform a binary-discrimination exercise where they matched a target image to one of the previously presented flanker stimuli (rules). The tasks included two types of trials: switch trials and non-switch or stay trials. During switch trials, the flankers to which the participants were discriminating the targets from changed i.e. there was a switch in the task rules. These events were indicated by the presentation of the warning red rectangles and fixation cross in a new location, followed by a new set of flankers. During non-switch trials, the task rules, i.e. the relevant flankers, remained unchanged and only a red fixation cross (500ms) preceded the appearance of a new target image.

The task started with a rest period (16s) indicated by the presence of a black fixation cross centred in the middle of a white screen. Next, two red rectangles and a red fixation cross were drawn to the screen for 500ms, which indicated the participants the location of the coming relevant stimuli. After this, either one (Study 1) or four (Study 2) sets of flanker images were presented for 4s. In Study 2, the red fixation cross remained presented in the screen to direct the participants to the relevant set of flankers. After four seconds, the flankers of interest (i.e. the two images and the fixation cross) were replaced by a target image centred were the fixation cross was. The participants were asked to select to which flanker the target was most similar to as fast as possible. 1500ms were given to the participants to respond with a left or right button press.

### 0.5 Task parameters and experiment space

The set of stimuli, created by [29], had four categories of images: male faces, abstract lines, abstract figures and rooms. Stimuli of varying degree of similarity were created by morphing within a category (example in Figure 10). For the task in Study 2, the target images were of 1° of similarity to one of the flanker stimuli.

**Figure 10.**
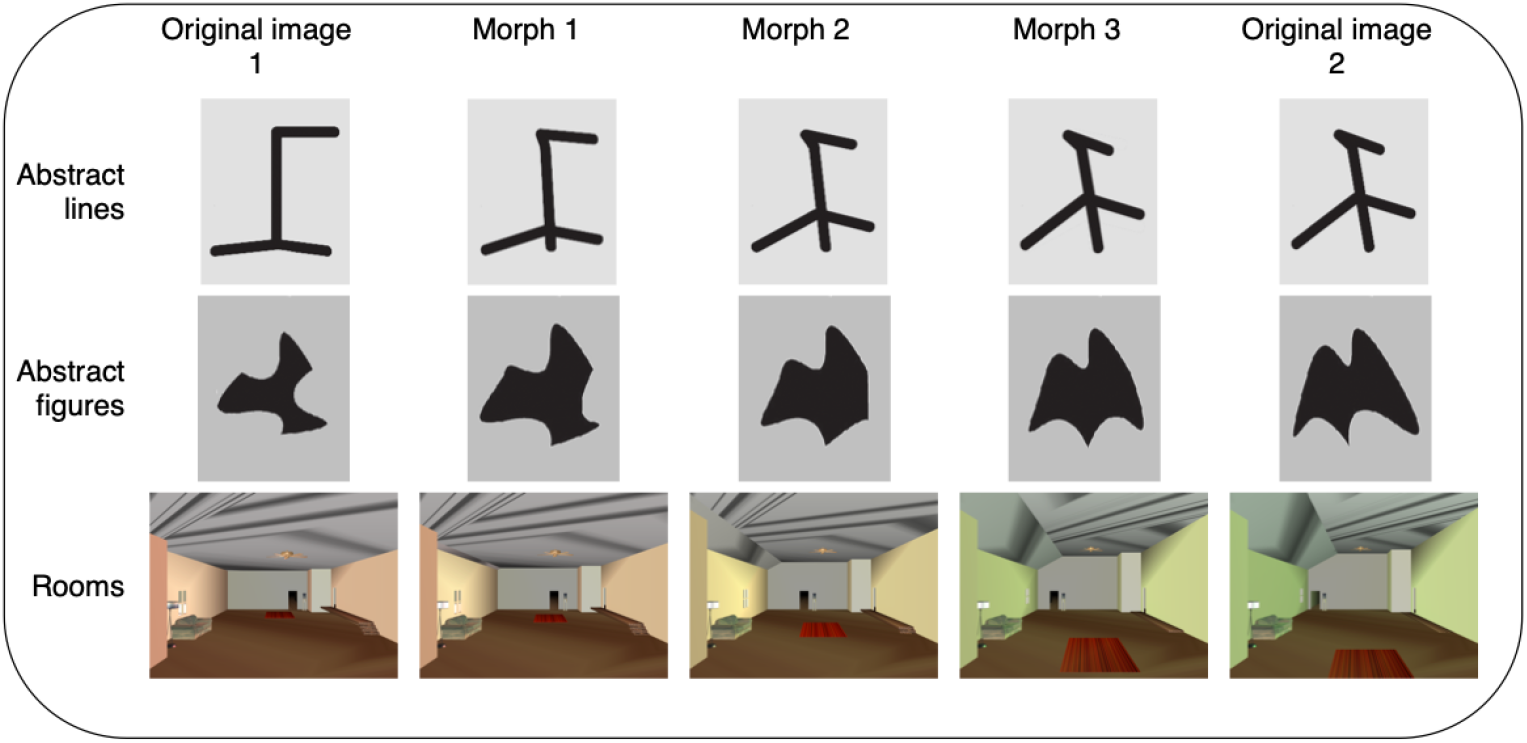
Stimuli set. Four categories of stimuli were used for the switching task: male faces, abstract lines, abstract figures and rooms. The original images were morphed with each other to create stimuli at various degrees of similarity. Faces are not displayed here to avoid potentially identifying information.

#### 0.5.1 Study 1

The complexity of the task was modified in 3 dimensions: switch distance, task discrimination and sequential predictability. The first dimension represented the similarity of the flankers between switch trials (switch distance) and it had three levels: switches of 1° similarity, switches of 3° similarity and category switch. The second dimension represented the similarity of the target to the flankers (discrimination) and it had 3 levels: same item, item at 1° similarity and item at 3° similarity. The third dimension represented the sequential predictability of switch trials (switch predictability) and it had 2 levels: switches at a regular point in the sequence, and switches at pseudo-random points in the sequence. The total number of switches remained the same for each sequence. Thus, a 3×3×2 experiment space was created.

From the experiment space, nine dimensional combinations were defined (Figure 1D), which were used as task parameters during the experiment. It should be noted that there were too many conditions to sample all of them using a traditional offline fMRI study design; instead, the selected combinations allowed the relevant dimensions to be examined in this initial study. Nonetheless, the sampling was sufficient to determine which dimensions appeared to have an effect on the activation of DMN regions.

#### 0.5.2 Study 2

The parameters of the cognitive paradigm were modified within the predictability dimension along three components: sequential, spatial and perceptual predictability of switch events. The sequential aspect referred to the frequency at which switch events occurred; the spatial component was linked to the location of the new relevant stimuli; and the perceptual aspect alluded to the nature of the incoming relevant stimuli i.e. the category of the rules (faces, abstract lines, abstract lines or rooms). The dimensions were chosen based on the nature of the cognitive task and limited to a number that allowed the experiment space to be sampled in a single session.

A 2×2×2 experiment space was constructed by assigning to each sub-dimension two levels of complexity. The switch events in the first level were characterised by having a highly predictable sequence: sequentially there was a change in rules every four trials, with every switch the location of the stimuli shifted one position downwards, and perceptually the stimuli changed from faces to abstract lines to rooms to abstract figures. On the other hand, the switches in the second level followed a pseudo-random sequence: sequentially a switch could occur at any trial within a grouping of four trials (keeping the total number of switches constant), spatially the relevant stimuli could switch one position upwards or downwards (maintaining the frequency of direction a maximum of two switches) and perceptually the category of the stimuli was randomly chosen (ensuring it did not follow the predictable sequence and it was distinct from the preceding rule).

The size of the experiment space allowed for a full-factorial experiment to be conducted, where every possible combination of dimensions was sampled (Figure 1E). For example, in a block the task could have sequential complexity of level 1, spatial complexity of level 2 and perceptual complexity of level 1.

### 0.6 Experimental procedure

The participants were subjected to three back-to-back fMRI runs. Within each run, the participants performed 9 blocks of 20 trials of the switching task (Study 1) or 8 blocks of 30 trials of the switching task (Study 2), with 16 seconds of rest in between. The parameters of each block corresponded to one of the previously defined combinations (Figure 1D-E), and the order of each block was randomised across runs and participants. Participants were trained in the task by performing one practice block outside the scanner before the experimental procedure. In the scanner, the task was projected onto a screen that the participants could see through mirrors placed on the head coil and the responses were recorded using a pair of response grips (ResponseGrip, NordicNeuroLab AS, Bergen, Norway).

### 0.7 fMRI acquisition

MRI data were acquired in a 3T Siemens Verio (Siemens, Erlangen, Germany) using a 32-channel head coil. T1-weighted structural images were acquired using an MP-RAGE sequence, isotropic voxel of 1mm3, repetition time (TR) of 2300ms, echo time (TE) of 2.8ms, inversion time of 900ms, flip angle (FA) of 90°, field of view of 256×256mm, 256×256mm matrix, 160 slices and GRAPPA acceleration factor of 2. For the fMRI images, a T2°-weighted echo-planar imaging (EPI) sequence was acquired using an isotropic voxel of 3mm3, TR of 2s, a TE of 30ms, FA of 80°, field of view of 192×192×105mm, 64×64 matrix, 35 slices and GRAPPA acceleration factor of 2.

### 0.8 fMRI pre-processing

The fMRI Expert Analysis Tool (FEAT, Version 6.00) from the FMRIB’s Software Library [FSL [34, 53]] was employed for the pre-processing and analysis of the fMRI data. FMRIB’s Linear Image Registration Tool [FLIRT [34, 35]] was used to register the extracted brain tissue [BET [52]] to the Montreal Neurological Institute (MNI) standard atlas. Motion-correction of the images was performed with MCFLIRT [33], and spatially smoothed with a 5mm Gaussian kernel filter. A temporal high-pass filter with a cut-off of 100s (Gaussian-weighted least-squares straight line fitting) was applied to remove low frequency artefacts. The EPI sequences were registered to the MNI space using the T1-weighted images as intermediate by first carrying boundary-based registration [BBR [24]] to the main structural image followed by affine registration to the standard brain space [34, 35].

### 0.9 fMRI task analysis

#### 0.9.1 Whole-brain analysis

FEAT was used to analyse the pre-processed fMRI data. General linear models (GLM) were constructed with regressors corresponding to task blocks (all trials included in a block) and switch trials (only trials where new rules were presented) for each of the conditions tested (the 9 and 8 combinations of dimensions). Thus, 18 (Study 1) and 16 (Study 2) regressors of interest were fitted into each subject-level GLM. The task regressors were modelled by convolving a double-gamma haemodynamic response function (HRF) with a boxcar kernel. Movement-related noise was accounted for by adding six motion regressors to the design matrix. The following contrasts of interest were generated: all task blocks (“Task blocks > Rest”) and all switch events (“Switch > Rest”). A second-level analysis using a fixed effects model was performed to estimate each subject’s mean response across the three runs. This was done by forcing the random effects variance to zero in FLAME [FMRIB’s Local Analysis of Mixed Effects [6,68]] for each subject-level contrast. The resulting contrast values and variances were fed into a third-level analysis to combine data from all participants for the relevant contrasts using FLAME 1, the FMRIB local analysis of mixed effects [6,68]. A Gaussian random-field based cluster inference (threshold of z > 3.1 and cluster-correction significance threshold of p < 0.05) was applied to threshold the final Z statistical images.

#### 0.9.2 Region of interest analysis

Voxel-wise group-level analysis were performed for the DMN to explore the patterns of activation across the different switching conditions. The “Switch > Rest” contrast was investigated to identify regions of the DMN that displayed significant activation. Regions of interest (ROIs) were created (5mm radius spheres) placed at the peak activation coordinates of each node (“fslmaths” command, binarized). Peak activation of regions not belonging to the DMN but surviving cluster correction were also employed to define ROIs. For each of the eight switch trial regressors, the mean activation within each ROI was extracted (“fslmeants” command) and compared using a repeated-measured analysis of variance (ANOVA).

#### 0.9.3 Predictability dimension modelling

In order to disambiguate the brain activation in response to each predictability dimension in Study 2, a second higher-order model was constructed using the second-level switch trial contrasts. A regressor was defined for each of the three predictability dimensions, and a nuisance regressor was added to model the task effect. Contrasts were created for each regressor of interest (“Sequentially predictable > Unpredictable”, “Perceptually predictable > Unpredictable” and “Spatially predictable > Unpredictable”. The higher-order analysis was performed using FLAME 1 [6,68]. The resulting Z statistical images were thresholded using Gaussian random-field cluster inference (initial voxel-level threshold of z > 3.1 and cluster-correction at p < 0.05).

#### 0.9.4 ROI analysis of predictability dimensions

To explore whether there were sub-threshold activity patterns, active clusters identified from the whole-brain analysis were used to define ROIs. Active clusters were identified from the group activation maps from the predictability dimension contrasts using the FSL command “cluster” (voxel threshold of z > 3.1, uncorrected). Brain masks were created for clusters containing more than 10 voxels using the FSL command “fslmaths”. The binarised masks were employed in the group-level dimension model, with the same predictors and contrasts (“Sequentially predictable > Unpredictable”, “Perceptually predictable > Unpredictable” and “Spatially predictable > Unpredictable”). The higher-order analysis was performed using FLAME 1 [6, 68]. The resulting Z statistical images were thresholded using Gaussian random-field cluster inference with cluster-correction at p < 0.05 at a voxel-level thresholds of z > 3.1. The mean activation within each cluster was extracted (“featquery” command) and compared using a t-test against zero with correction for multiple comparisons with Bonferroni.

### 0.10 Statistical analysis

Data analyses were performed using [41] and [59]. Mean accuracy, defined as the percentage of correct responses, and mean reaction times for switch and stay trials were computed and analysed using ANOVA. Significance was set at p < 0.05 and post hoc tests were performed with Tukey correction to correct for multiple comparisons unless otherwise stated.

## Acknowledgments

We thank Ivana Russo for helping in data acquisition and all the participants that took part in the study for their time. We are grateful to Dr. Stefano Sandrone for his input on previous versions of the manuscript.

